# Loss of *Lkb1* cooperates with *Braf*^*V600E*^ and UV radiation increasing melanoma multiplicity and neural-like dedifferentiation

**DOI:** 10.1101/2024.03.29.587140

**Authors:** Kimberley McGrail, Elena González-Sánchez, Paula Granado-Martínez, Roberto Orsenigo, Yuxin Ding, Berta Ferrer, Javier Hernández-Losa, Iván Ortega, Juan M. Caballero, Eva Muñoz-Couselo, Vicenç García-Patos, Juan A. Recio

## Abstract

The mechanisms cooperating with *BRAF*^*V600E*^ oncogene in addition to ultraviolet (UV) radiation in melanoma development are of great interest. Analysis of human melanoma tumors (TCGA) indicates that 50% or more of the samples express no or low amounts of LKB1 protein. Here, we report that the concomitant neonatal *Braf*^*V600E*^ activation and *Lkb1* tumor suppressor ablation in melanocytes led to full melanoma development. A postnatal single-dose of UVB radiation had no effect on melanoma onset in *Lkb1*-depleted mice in respect to *Braf*^*V600E*^-irradiated mice, but increased tumor multiplicity. In agreement to this and previous reports, *Lkb1* null irradiated mice showed a deficient DNA damage repair (DDR). Histologically, tumors lacking *Lkb1* were enriched in neural-like tumor morphology. Genetic profiling and gene set enrichment analyses of tumor samples-mutated genes indicated that loss of *Lkb1* promoted the selection of altered genes associated to neural differentiation processes. Thus, these results suggest that loss of *Lkb1* cooperates with *Braf*^*V600E*^ and UVR impairing DDR and increasing melanoma multiplicity and neural-like dedifferentiation.

## INTRODUCTION

Cutaneous melanoma is the most aggressive form of skin cancer, responsible for the majority of skin cancer related deaths. In melanoma, *BRAF* mutation is the most frequent alteration (50% of melanoma tumors) leading to abnormal activation of the RAS/RAF/MEK/ERK signaling pathway (Davies et al., 2002; Greaves et al., 2013; Long et al., 2011). There are no differences between *BRAF* mutation frequency of benign and tumoral lesions (Pollock et al., 2003), indicating that mutated *BRAF* is not sufficient for melanoma transformation (Michaloglou et al., 2005). Multiple studies have identified major host and environmental risk factors for melanoma. The predominant environmental risk factor is exposure to UV radiation (UVR), responsible of the generation of DNA mutations that, if not properly repaired, can result in genomic instability and consequently, in tumorigenesis (Armstrong and Kricker, 2001; Noonan et al., 2001). The cooperation of UVR and *BRAF* mutations promoting malignant melanoma has been previously described under different contexts (Luo et al., 2013; Viros et al., 2014). In addition to UVR, expression of BRAF^V600E^ combined with silencing of the tumor suppressor PTEN (Dankort et al., 2009), or the downregulation of the tumor suppressor NF1 (Gibney and Smalley, 2013), induced melanoma development and progression.

Serine threonine kinase 11 (*STK11*), also known as LKB1, is a ubiquitously expressed and evolutionary conserved serine/threonine kinase identified as a tumor suppressor. It is involved in a number of biological processes, including DNA damage repair (DDR) (Esteve-Puig et al., 2014; Gupta et al., 2015; Ui et al., 2014; Wang et al., 2016). *LKB1* has been found mutated in several tumor types, like lung cancer (Ji et al., 2007; Sanchez-Cespedes et al., 2002), cervical cancer (McCabe et al., 2010), pancreatic cancer (Morton et al., 2010; Su et al., 1999), breast cancer (Shen et al., 2002) and malignant melanoma (Guldberg et al., 1999; Rowan et al., 1999). In fact, Peutz-Jeghers syndrome (PJS), which results from germline mutations in *LKB1*, is characterized by an increased probability of developing cancer (Giardiello et al., 2000; Lim et al., 2004). There are several studies highlighting that cancer development and progression can be promoted in the absence of *LKB1* even in haploinsufficiency (Esteve-Puig et al., 2014). In the case of melanoma, it has been described that *Lkb1* inactivation facilitates the expansion of melanoma pro-metastatic tumor cell subpopulations upon RAS pathway activation (Liu et al., 2012). In addition to this, LKB1 inactivation synergized with *BRAF*^*V600E*^ promoting cell invasion and migration (Zhang et al., 2018), and prevented *BRAF*^*V600E*^ oncogene-induced growth arrest (Damsky et al., 2015). Moreover, *Lkb1* deficiency sensitized mice to DMBA-induced skin and lung squamous cell carcinomas (Gurumurthy et al., 2008) and increased the progression of lung adenomas into carcinomas (Gonzalez-Sanchez et al., 2013). Additionally, LKB1 plays a crucial role in the regulation of neural crest cell migration and differentiation during embryonic development, where dysregulation of LKB1 function can lead to defects in neural crest cell development and contribute to various developmental abnormalities (Radu et al., 2019).

It is known that melanoma tumors are characterized by a high mutational burden, resulting from exposition to UVR. *LKB1* loss contributes to tumor development and progression in several tumor types under different genetic contexts, including *BRAF* mutant melanoma. Knowing that LKB1 has a relevant role in DDR (Esteve-Puig et al., 2014) and neural crest cell differentiation (Radu et al., 2019), we investigated the contributions of *Lkb1* loss to UVR-induced melanoma development and progression in a *Braf*^*V600E*^-mutant animal model. Our data show that a substantial fraction of *BRAF*-mutated human melanomas either do not express or express low amount of LKB1. *In vivo* modeling of UV-induced *Braf*^*V600E*^-mutated melanoma showed that loss of *Lkb1* cooperates with *Braf*^*V600E*^ and UVR impairing DDR and increasing melanoma multiplicity and dedifferentiation.

## MATERIAL AND METHODS

### Mouse Model

All mice were cared for and maintained in accordance with animal welfare regulations under an approved protocol by the Institutional Animal Care and Use Committee at Vall d’Hebron Institut de Recerca (VHIR) and Biomedical Research Park of Barcelona (PRBB). The *Tyr::Cre*^*ERT2*^*;Braf*^*CA*/CA^ strain (Dankort et al., 2007) and the *Tyr::Cre*^*ERT2*^;*Braf*^*CA/CA*^;*Lkb1*^*F/F*^ strain (González-Sánchez et al., 2013) were previously described. Both sexes were used for experiments.

#### Melanoma development in vivo

Mice were treated topically on postnatal day 2.5 and 3.5 with 100 μl of an acetone solution containing 5 mg/ml 4-hydroxytamoxifen (4OHT) (Sigma, St Louis, MO, USA). Neonatal mice were irradiated on postnatal day 3.5 as previously described (Recio et al., 2002). Data for tumor development was analyzed by Kaplan-Meier survival analysis.

### Human melanoma samples

All pseudonymized human melanoma samples were provided by the Vall d’Hebron Research Institute under the National Research Ethics Service approved study number PR(AG)59-2009. Informed consent was provided by all patients.

### Immunohistochemistry (IHC)

For IHC, 4 μm sections of formalin-fixed paraffin-embedded skin or tumor samples were stained following the manufacturer’s antibody protocol. The samples were developed by using either secondary antibodies linked to horseradish peroxidase with the UltraView™ Universal DAB Detection Kit (Ventana Medical System, Roche) or secondary antibodies linked to fluorophores. Staining was performed either manually or using the automated immunostainer Beckmarck XT (Ventana Medical Systems). For the samples processed manually, antigen retrieval was performed using target retrieval solution pH 6.0 or pH 9.0 according to antibody protocol recommendations (Agilent, Santa Clara, CA, USA). The samples were scanned (panoramic slide digital scanner) and evaluated by two independent pathologists (using 3DHistech Software). LKB1 (#13031; 1:250), p-AKT^S473^ (#4060; 1:100), p-S6^S235/236^ (#2211; 1:500) and p-4EBP1^T37/46^ (#2855; 1:1000) antibodies were obtained from Cell Signaling (Danvers, MA, USA). Cre antibody (NB100-56135; 1:1000) was obtained from Novus Biologicals (Littleton, CO, USA). BRAF^V600E^ (#760-5095) was purchased from Ventana Medical System. TRP2 (Dct, Pep8) was obtained from Dr. Vincent Hearing (National Institute of Health) (Tosti et al., 1994). Secondary fluorescent antibodies were purchased from Thermo Fisher Scientific (Waltham, MA, USA).

### Dot Blot

Back skins of *Tyr::Cre*^*ERT2*^*;Braf*^*CA/CA*^*;Lkb1*^*F/F*^ untreated and 4OHT-treated 2.5 days-neonates were collected 20 hours and 7 days after treatment with UVR. Genomic DNA was isolated using the DNeasy kit (Qiagen, Venlo, Netherlands) following the manufacturer’s recommendations. A total amount of 100 ng of DNA was resuspended in 0.5M NaOH; 10 mM EDTA, denatured and spotted on a nitrocellulose membrane using the Bio-Dot SF (Bio-Rad, Hercules, CA, USA). Then, the membrane was heated at 80 ºC for 2 hours and incubated with primary antibodies against 6-4 photoproducts (6-4 pps), Dewar photoproducts (Dewar pps) and cyclobutane pyrimidine dimers (CPDs), purchased from CosmioBio (Carlsbad, CA, USA). Secondary antibodies were obtained from GE Healthcare (Chicago, IL, USA). Bound antibodies were detected by ECL (GE Healthcare). DNA loading was assessed by re-probing the membrane with radiolabeled mouse genomic DNA with ethidium bromide (Sigma). Quantification of the spots was performed using ImageJ 1.53a.

### Whole-exome sequencing (WES)

WES and data analysis were performed at the Genomic Facility of VHIO (Vall d’Hebron Oncology Institute). In brief, DNA genomic libraries were prepared from fresh tissue (tumor and nontumor) and FFPE tumor tissue DNA prior to exome capture (SureSelect XT Human All Exon v5, Agilent). Libraries were sequenced on a HiSeq2000 instrument (Illumina) with a mean coverage of 100X. Reads were aligned, and somatic variants called by comparison with nontumor samples (VarScan2).

### Data availability

The gene alteration data used in this study are publicly available at cBioPortal (Skin Cutaneous Melanoma; TCGA, PanCancer Atlas, 448 samples; https://www.cbioportal.org). The exome sequencing datasets reported in this article have been deposited in the National Center for Biotechnology Information (NIH) under accession number PRJNA1073850.

### Statistics

Statistical analyses were performed in GraphPad Prism 9.0 (GraphPad Software Inc.) using a 2-tailed Student’s t test to compare differences between two groups.

## RESULTS

### A substantial fraction of *BRAF*^*V600E*^ mutant human melanomas do not express LKB1

It is known that loss of *PTEN* and *NF1* tumor suppressors, as well as UVR, cooperate with *BRAF*^*V600E*^ in melanoma development and progression (Dankort et al., 2009; Gibney and Smalley, 2013). In the case of *Lkb1* it was suggested that its inactivation abrogated *Braf*^*V600E*^-induced cell growth arrest (Damsky et al., 2015). Furthermore, loss of *Lkb1* impaired UV-induced DDR leading to genetic instability and the development of skin tumors (Esteve-Puig et al., 2014). To further analyze the role of the tumor suppressor LKB1 in melanomas carrying the *BRAF*^*V600E*^ mutation, we analyzed the expression and mutational status of *LKB1* from 448 human samples obtained from the TCGA database (Skin Cutaneous Melanoma; TCGA, PanCancer Atlas). The analysis of the data showed a tendency for the negative correlation between *LKB1 (STK11)* and *BRAF* mRNA expression, where the absolute numbers of *LKB1* transcripts were more abundant than the *BRAF* mRNA molecules (Figure 1A). Contrary to expected, most tumors expressed low amounts of LKB1 while BRAF protein was highly expressed, including *BRAF-*mutated samples (Figure 1A). The correlation between the respective amounts of *LKB1 or BRAF* mRNA and protein supported this observation (Figure 1B). In agreement to this, the analysis of the putative copy number of *LKB1* and *BRAF* genes showed that *LKB1* genetic modifications were related to shallow deletions, while *BRAF* genetic alterations principally corresponded to gains and amplifications, including *BRAF*-mutated samples (Figure 1C). We validated these results by immunohistochemistry in an independent set of human samples (14 human *BRAF*^*V600E*^-mutated melanomas). H-Score quantification of the samples showed that while 85% of the samples showed a positive staining for BRAF^V600E^, 42% of the samples showed either low or no expression of LKB1 protein (Figure 1D). Altogether, these results suggest that a substantial fraction of *BRAF*^*V600E*^-mutated melanomas might have LKB1-dependent functional impairments.

**Figure 1:**
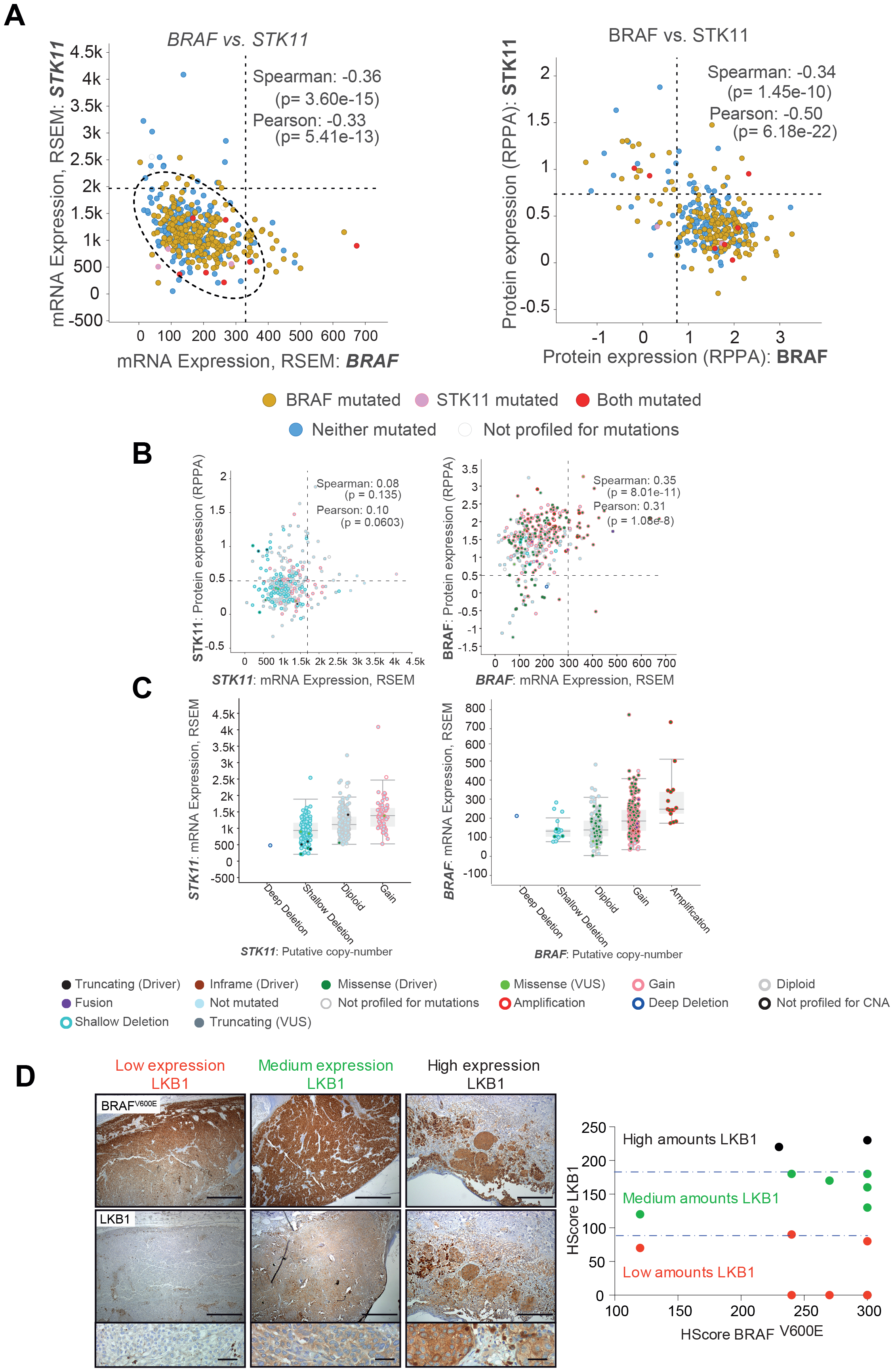
Most BRAF^V600E^ mutant human melanomas express low or absent amounts of LKB1. **(A)** On the left, correlation between *BRAF* and *STK11* mRNAs in human melanoma samples. On the right, correlation between BRAF and LKB1 protein expression in human melanoma samples. Data obtained from the cBioPortal database (Skin Cutaneous Melanoma; TCGA, PanCancer Atlas). **(B)** On the left, correlation between the expression of *STK11* mRNA and the expressed LKB1 protein. On the right, correlation between the expression of *BRAF* mRNA and the expressed BRAF protein. Data obtained from the cBioPortal database (Skin Cutaneous Melanoma; TCGA, PanCancer Atlas). **(C)** Analysis of putative *STK11* and *BRAF* genomic copy numbers in relation to mRNA expression. Data obtained from the cBioPortal database (Skin Cutaneous Melanoma; TCGA, PanCancer Atlas). **(D)** LKB1 immunostaining of BRAF^V600E^ human melanomas. Representative pictures of tumors expressing low, medium and high LKB1 amounts are shown. On the right, the graphic representation of LKB1 HScore against BRAF^V600E^ HScore is shown.

### Neonatal loss of *Lkb1* cooperates with *Braf*^*V600E*^ in melanoma development and increases tumor multiplicity in UVR induced melanomas

Epidemiological studies have demonstrated a strong association between UV radiation and melanoma risk, where DNA is the major target of direct or indirect UV-induced cellular damage. Due to the LKB1 role in DDR (Esteve-Puig et al., 2014) and the above observation suggesting the lack or low expression of LKB1 in *BRAF*^*V600E*^ mutant tumors, we investigated the contribution/s of *Lkb1* loss to UVR-induced melanoma development and progression in a *Braf*^*V600E*^ mutational context. To that end, we used the conditional and inducible Tyr::*Cre*^ERT2^;*Braf*^*CA*^ mice (Dankort et al., 2009). *Braf*^CA^ mice express wild type BRAF prior to Cre-mediated recombination upon 4OH-tamoxifen treatment, at which time oncogenic *Braf*^V600E^ is expressed in physiological amount. To generate the *Tyr*::*Cre*^ERT2^;*Braf*^CA^; *Lkb1*^F/F^ mouse we crossed the *Tyr*::*Cre*^ERT2^;*Braf*^CA^ mouse with the conditional knockout *Lkb1*^flox/flox^ (*Lkb1*^*F*/F^) mouse. As previously described, activation of BRAF did not promote melanoma development except for one old homozygous *Braf*^*CA/CA*^ mice. However, UVB irradiation promoted melanoma development in 85.7% of *Braf*^CA/+^ and 87.5% of *Braf*^CA/CA^ mice. In contrast to previously suggested (Damsky et al., 2015), *Lkb1* haploinsufficiency cooperated for melanoma development with the activation of either one *Braf*^CA/+^ allele (18.1%) or both *Braf*^CA/CA^ alleles (31.2%) albeit at late time points (mean onset of 399±38.9 and 285±45 days, respectively). However, the loss of both *Lkb1* alleles in mice carrying either one *Braf* mutant allele (*Braf*^CA/+^) or both *Braf* mutant alleles (*Braf*^CA/CA^) promoted a slight decrease in the incidence of melanoma from 18.1% to 11% and from 31.2% to 12.5%, with an onset of 326±90 and 319±17 days, respectively (Figure 2A). In comparison with UVR-induced melanomas, the tumor incidence was not further increased upon *Lkb1* lost in either *Braf*^CA/+^ or *Braf*^CA/CA^ mice (Figure 2B). Nevertheless, the loss of the second *Lkb1* allele promoted a slight delay in melanoma development together with an increase in tumor multiplicity (Figure 2C, D). Overall, the above results showed that loss of *Lkb1* cooperates with *Braf*^*V600E*^ in melanocyte transformation allowing melanoma development. Furthermore, loss of *Lkb1* increased tumor multiplicity, particularly in response to UVR.

**Figure 2:**
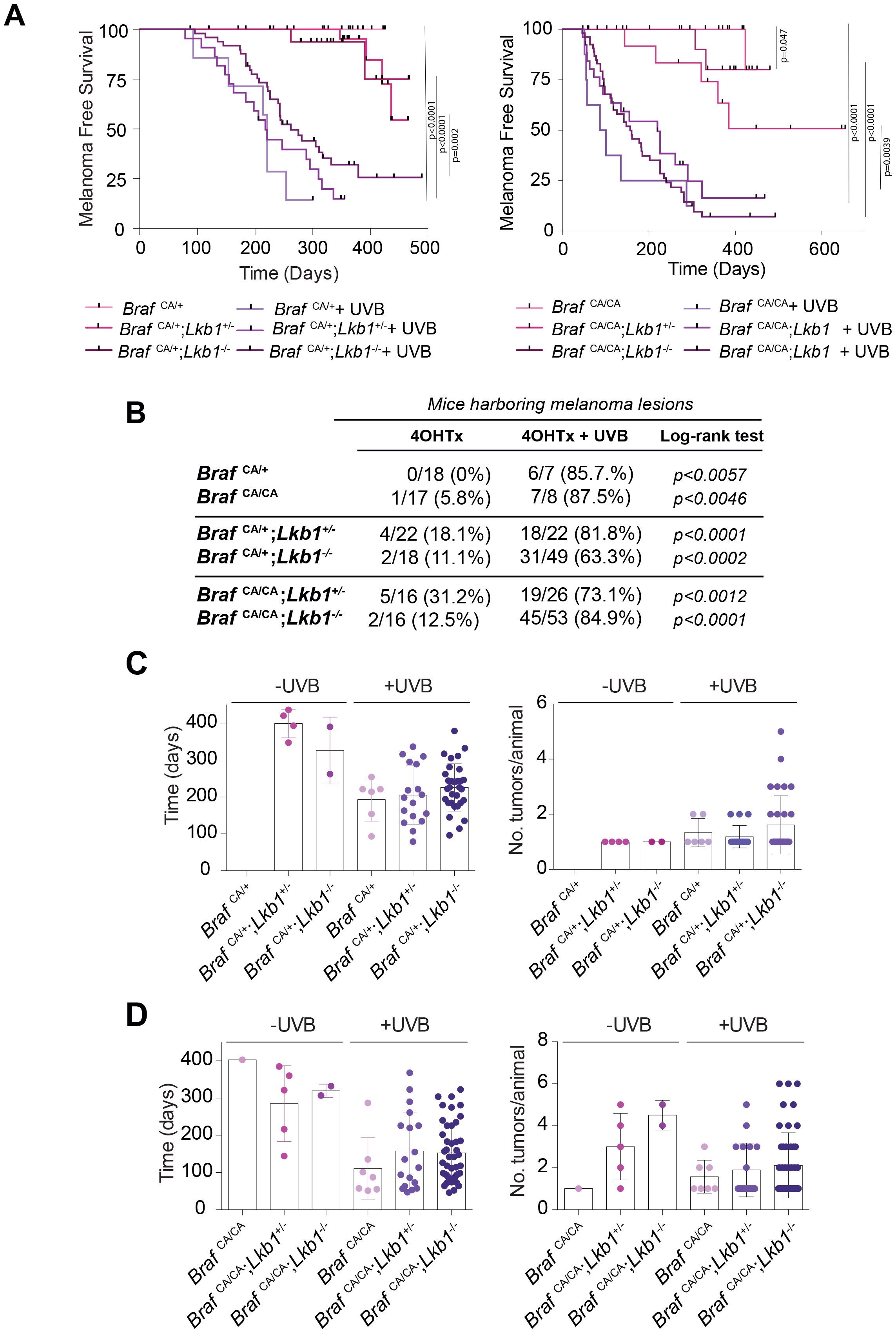
Loss of *Lkb1* cooperates with *Braf*^*V600E*^ mutation promoting melanoma development. **(A)** Kaplan-Meier survival curve of mice treated with 4OHTx or 4OHTx + UVR. *Braf*^*CA/+*^ *(n=18) / Braf*^*CA/+*^; *Lkb1*^*+/-*^ *(n=22) / Braf*^*CA/+*^; *Lkb1*^*-/-*^ *(n=18) / Braf*^*CA/CA*^ *(n=17) / Braf*^*CA/CA*^; *Lkb1*^*+/-*^ *(n=16) / Braf*^*CA/CA*^; *Lkb1*^*-/-*^ *(n=16)*. **(B)** Table showing the total number of mice per group and the percentage of mice developing melanoma tumors in response to either 4OHTx or 4OHTx + UVR treatments. p-values for the calculated incidences are indicated. **(C)** Mean tumor onset and tumor multiplicity of *Braf*^*CA/+*^ mice with or without one or both copies of *Lkb1*. p-value was calculated by Student’s t-test. **(D)** Mean tumor onset and tumor multiplicity of *Braf*^*CA/CA*^ mice with or without one or both copies of *Lkb1*. p-value was calculated by Student’s t-test.

### UVR- and *Lkb1*^F/F^-induced melanoma tumors showed activation of mTORC1/2

It has been previously described that *in vivo Braf*^*V600E*^-induced melanoma models require the concomitant activation of both mTORC1 and mTORC2/Akt for cell progression to malignancy (Damsky et al., 2015).

The same study suggested that *Lkb1* loss abrogated *Braf*^*V600E*^-induced cell cycle arrest, however, did not lead to melanoma formation. Our data showed that loss of *Lkb1* (even in haploinsufficiency) allowed the development of *Braf*^*V600E*^ mutant melanomas with a low incidence (18%-30%) 10 months after *Braf*^*V600E*^ activation and *Lkb1* knock out, most likely due to an increase in genomic instability. Since *Lkb1* inactivation will lead to mTORC1 activation, we investigated whether these tumors acquired also the activation of mTORC2. Independently of the UV irradiation and *Lkb1* status, all analyzed tumors stained positive for surrogate markers of mTORC1 activation, p-4EBP1^T37/46^, p-S6^S235/236^ and mTORC2-mediated phosphorylation of pAKT^S473^ (Figure 3), confirming that activation of both mTORC1/2 is required for full *BRAF*^*V600E*^-induced melanomagenesis. Thus, these results support that loss of *Lkb1* abrogates *Braf*^*V600E*^-induced cell growth arrest allowing the malignant transformation over time and the activation of mTORC1/2.

**Figure 3:**
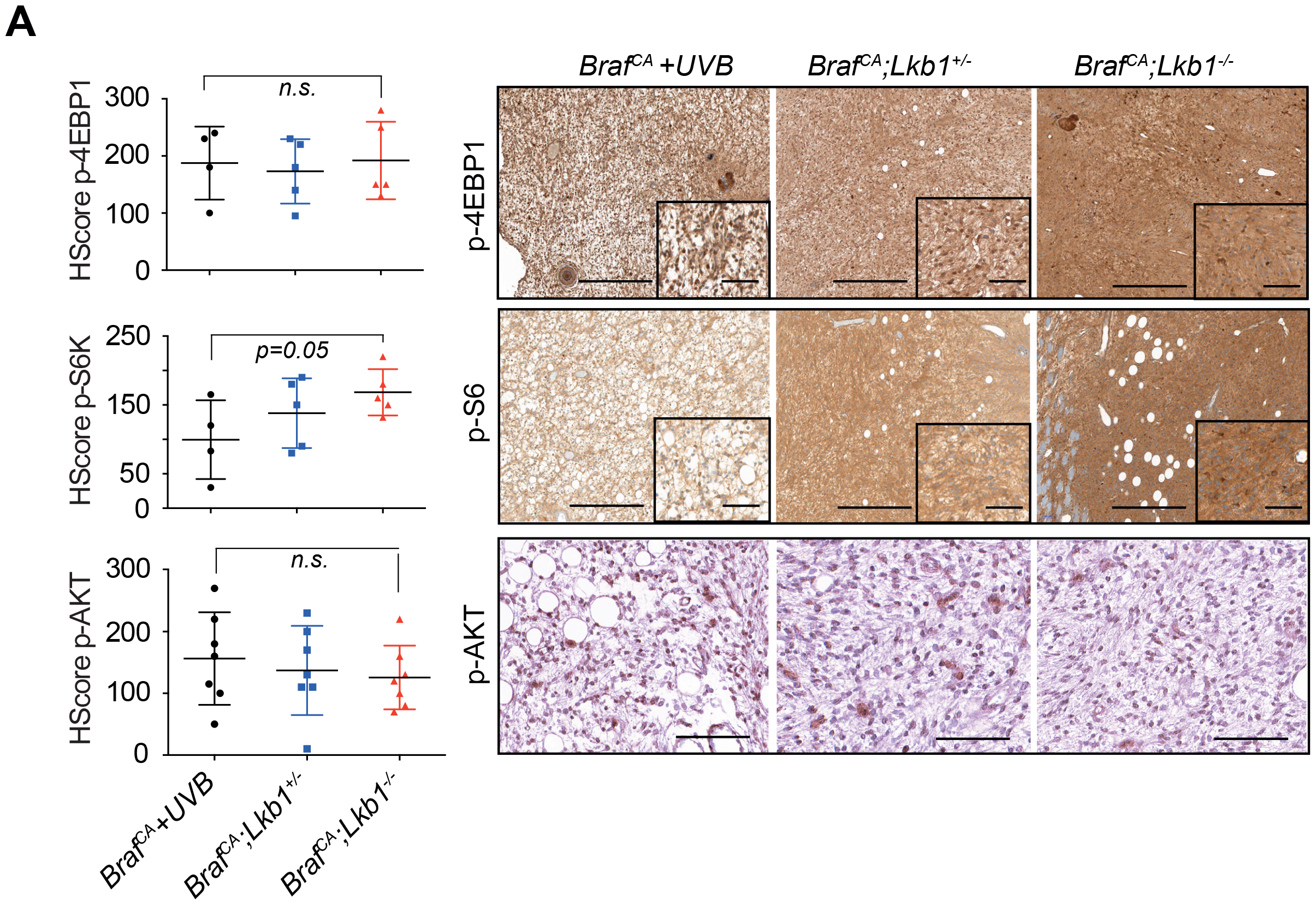
Activation of both mTORC1 and mTORC2 is necessary for *Lkb1* loss-driven melanoma progression. Representative immunohistochemistry images of tumors aroused in mice with the indicated genetic backgrounds showing the staining for p-4EBP1^T37/46^, p-S6^S235/236^ and p-AKT^S473^. Graphs on the left show the HScore for these markers in n=7 different tumors from each genetic background. p-value was calculated by Student’s t-test. Scale bars corresponds to 500μm.

### Loss of *Lkb1* increases tumor heterogeneity and impairs UVR-induced DDR

As previously described (Damsky et al., 2015; Dankort et al., 2009; Viros et al., 2014) melanomas were mainly amelanotic, localized in the dermis or subcutaneous, with no junctional component and occasionally displaying several types of morphologies. We distinguished three major tumor morphologies: melanoma with myxoid features; spindle-shaped to round plump cells melanomas, and melanomas with neural differentiation (Figure 4A). *Braf*^*V600E*^ mutant tumors derived from UV radiation or *Lkb1* loss were phenotypically heterogeneous (Figure 4A), nevertheless we could correlate the predominant type of lesions according to the mouse genotype and treatment (UVR). Melanomas with myxoid like morphology and spindle melanomas were predominant, and they developed mostly in response to UVR and independently of the absence of *Lkb1*. However, melanomas with neural differentiation were more frequent when *Lkb1* was deleted, with or without UVR, suggesting the existence of tumor morphologies preferentially linked to *Lkb1* dysfunction (Figure 4B, left). We analyzed the number of the different tumor morphologies according to the genetic dose of mutant *Braf*, alone and in combination with the loss of one or both *Lkb1* alleles. Overall, the loss of *Lkb1* increased tumor heterogeneity. UV-independent myxoid-like melanomas were absent in either *Braf*^CA/+^; *Lkb1*^-/-^ or *Braf*^CA/CA^; *Lkb1*^-/-^ and UV-independent melanomas with neural differentiation were mainly present in mice harboring the activation of two alleles of *Braf* (*Braf*^CA/CA^) and the deletion of either one or both alleles of *Lkb1* (*Lkb1*^*+/-*^; *Lkb1*^*-/-*^) (Figure 4B, right). Due to the observed tumor cells phenotypic heterogeneity and the predominance of certain histological subtypes depending on *Lkb1* expression status, we analyzed the expression of LKB1 in mouse melanocytes. Interestingly, co-staining of normal mouse skin (7-8 days postnatal) for LKB1 and the melanocyte marker TRP2 showed that only a small fraction of the mouse hair follicle melanocytes stained positive for LKB1 (Figure 4C), suggesting a possible connection between the tumor neural morphology and its cellular origin. Loss of *Lkb1* increased tumor heterogeneity. It is known that *Lkb1* loss impairs UVR-induced DDR generating genetic instability, sowe analyze the presence of UVB-induced DNA damage in mouse skins 20 hours and 7 days after UVB irradiation. 6-4 photoproducts (6-4pps), Dewar derivatives and cyclobutane pyrimidine dimers were still detected at high rates in *Braf*^CA^;*Lkb1*^-/-^ mice 7 days after UV irradiation, while it was almost totally repaired in wild type mice (Figure 4D) supporting the generation of genomic instability and consequently tumor heterogeneity.

**Figure 4:**
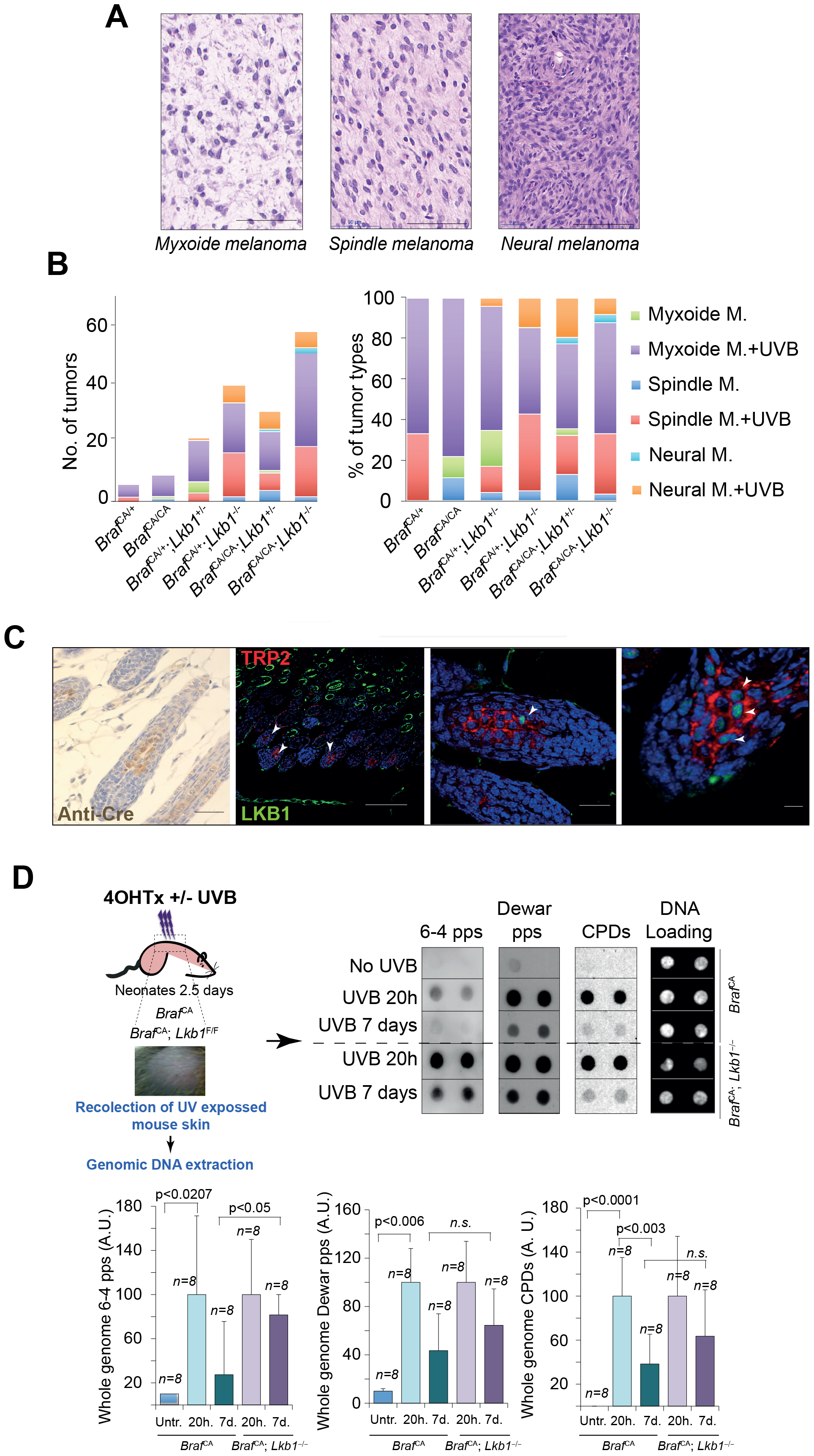
*Lkb1* loss promotes genomic instability and tumor heterogeneity. **(A)** Representative pictures of hematoxylin/eosin-stained tumors showing the histological melanoma subtypes. Bars represent 500μm **(B)** Graphs representing the number and percentage of the different melanoma subtypes according to the mouse genotype and the administered treatment. **(C)** Representative pictures of mouse skins stained with anti-Cre, LKB1 (green) and the melanocyte marker TRP2 (red). White arrowheads indicate melanocytes expressing LKB1. Scale bars correspond to 500 μm and 50 μm. **(D)** Immune Southern Blot (identification of DNA modifications by antibodies) of DNA samples from mouse skin 20 hours and 7 days after the indicated treatments. Graphs show the quantification of the generated DNA damage. The antibody signal obtained from the different assayed antibodies was normalized with respect to the total DNA amount. Error bars represent SEM (n=8). n.s. indicates not statistically significant differences.

### Tumor genetic profiling associated *Lkb1* loss to neural differentiation

To obtain insight about the mutations contributing to the melanocytic transformation by *Lkb1* loss and UVB-induced melanomagenesis, we performed whole exome sequencing in eight tumors generated in different genetic backgrounds (three spontaneous (one *Braf*^CA/CA^ (thereafter *B*) and two *Braf*^CA/+^;*Lkb1*^Δ/Δ^ (thereafter *B;L*)) and 5 UVB-induced tumors (three *Braf*^CA/+^ and two *Braf*^CA/+^;*Lkb1*^Δ/Δ^). We identified 1149 unique mutated genes among all analyzed mouse tumor samples (variant allele frequency; VAF>10%; Supplementary table 1). Analysis of the cutaneous melanoma TCGA database (PanCancer) showed 966 unique genes mutated in human samples (VAF>10%; Supplementary table 1). One hundred and seventy-seven of these mutated genes were identified in mouse and human melanomas, including *Braf* and *Stk11*. Gene set enrichment analysis showed that these common mutated genes were associated to tumor dysregulated processes including neural development (dedifferentiation), immune regulation, adhesion, motility and signal transduction (Figure 5A, Supplementary table 2). Mutational landscape analysis showed that C>T transitions were the most frequent nucleotide substitutions in *B*+UVB and *B;L*+UVB tumors (62.8% and 68.6%, respectively), while G>T transversions occurred more frequently in *B;L* tumors (77.5%) (Figure 5B). All tumor types exhibited similar frequencies in the types of alterations (the most frequent the non-synonymous variants), except for the stop gains, that were more frequent in the UVB induced melanomas (Figure 5C). Mutations targeted various protein family subtypes at different frequencies. *B;L*+UVB tumors showed an increased mutation frequency in protein kinases and transcription factors (Figure 5D). Then, we investigated which processes were affected by the identified mutated genes according to the genotype and treatment. Despite the low number of tumors per group of analysis, genes mutated in UVB irradiated tumors (VAF>20%) were significantly associated to cancer-related biological processes such as: extracellular matrix organization, adhesion, motility, (*Itga2b, Thbs4, Tnc, Itga1, Col6a5, Serpine1, Ptpn11, Cib1 and Ift74*), including the activation of RHO signaling (*Lmnb1, Myh9, Sh3bp1, Tuba3a, Zap70, Tiam2, Racgap1, Ndufs3, Rhobtb1, Dnmbp, Arhgef6, Dock8, Cenpi, Dock10, Fam91a1, Pde5a, Stard13*, and *Iqgap2*) involved in cytoskeletal dynamics, cell movement and associated to mTORC2 activation (Li and Gao, 2014; Senoo et al., 2019). Gene set enrichment analysis of the mutated genes in non-UVB irradiated *Lkb1* deficient tumors revealed their association to processes related to neural-like dedifferentiation (*Neurod6, Cttn, Dmd, Mark2, Hsp90ab1, Kif5a, Lamb2, Matn2, Thbs4, Tulp1, Gprin1, Cntnap1, Camsap2, Nrn1, Actl6b, Bbs4, Unc5b, Mcf2, Ntng2, Rnf165, Camsap1, Lrp4, Szt2, Gdf7, Nes, Plec*, and *Fat4*), adhesion, motility and also the dysregulation of RHO signaling pathway (*Atp6ap1, Cdc42ep3, Arhgap31, Mcf2, Fmnl3, Plekhg6, Ptpn13, Pkp4, Armcx3*, and *Baiap2l2*) (Figure 5E). Additionally, oncogene gene set analysis also showed that *B;L* tumors harbored mutation in genes associated to p53 (*Crim1, Mark2, Dmd, Amb2, Unc13b, Msln, Psmb8*), mTOR (*Adgre5, Cldn14, Irf7, Itpka, Alox15*, and *Ptprd*) and PTEN (*Fst, Nrn1, Pdzk1, Cldn14, Slc6a*, Slc26a4, *Tgfbr3, Kcnh7*, and *Lamc3)* activity. Identified mutated genes in UVB-irradiated *B;L* tumors were significantly involved in neural differentiation (i.e.: *Col3a1, Col5a1, Enah, Ncam1, Sema4d, Dpysl3, Arhgef7, Sptbn4, Ptk2b, Trpc5, Rorb, Nyap*, and *Pak4*) having also altered genes participating in signal transduction (*Egr4, Grid2, Fosb, Dyrk1b, Ror2, Prdm1, Ptk2b, Mapk10, Csf3r, Hras, Stat4, Six4, Rgs14, Grap, Pak4 Wnk2, Mapkbp1*, and *Arhgef7)* and immune modulating processes (*Cav1, Ifnb1 Stat4*, and *Il23r*) (Figure 5E). These results were also supported by the percentage of mutated genes belonging to the different signatures defining the melanoma cell subtypes (Tsoi et al., 2018) (Figure 5F and Supplementary table 2). Altogether, these results support the cooperation of *Lkb1* loss with *Braf*^*V600E*^ and its association to neural-like dedifferentiation.

**Figure 5:**
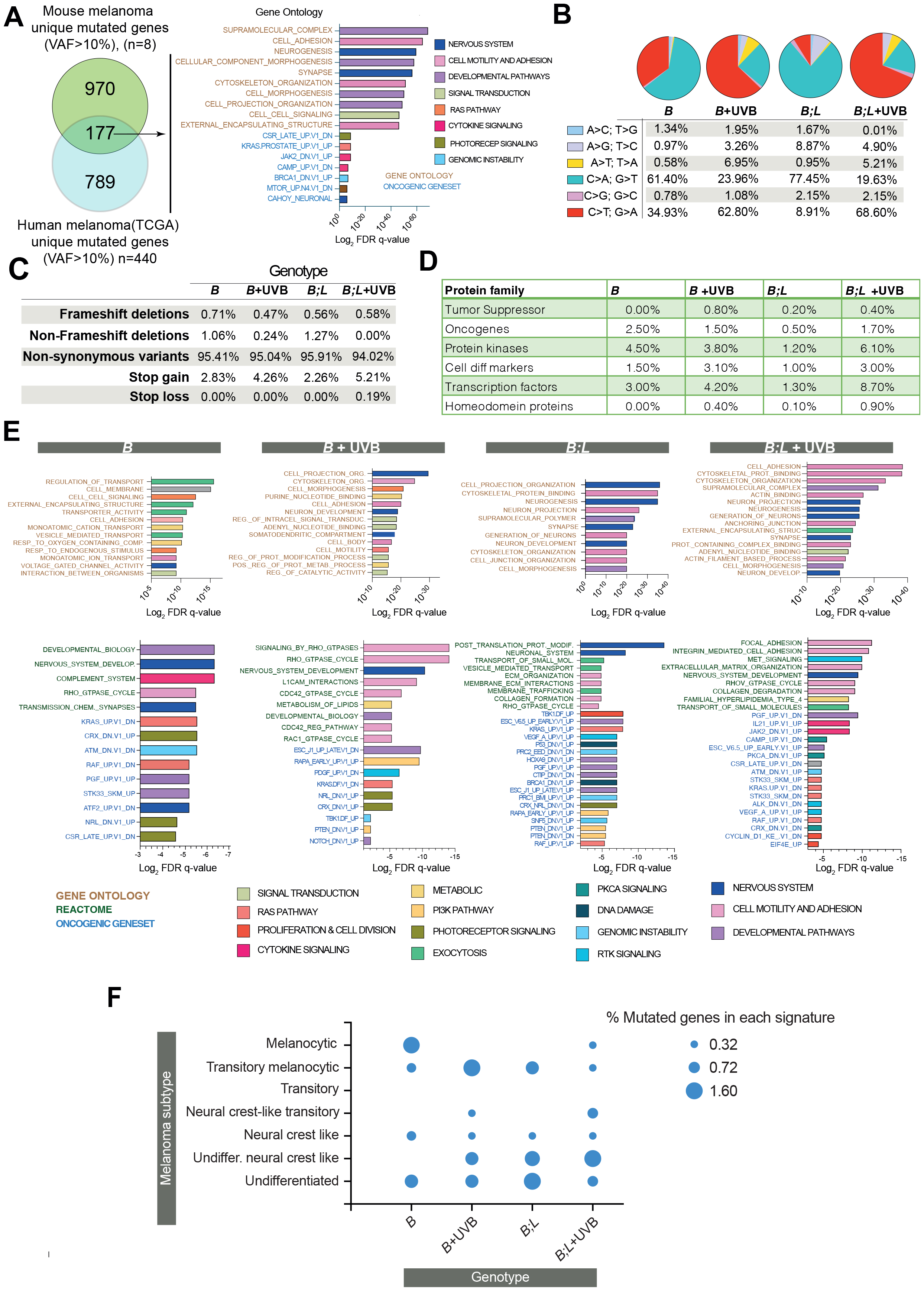
Exome sequencing mutational analysis. **(A)** Percentage of transitions and transversions detected in each genotype. **(B)** Percentage of different types of genetic alterations found in the indicated genotypes and treatments. **(C)** Percentage of mutated genes belonging to the indicated groups of proteins. **(D)** Gene set enrichment analysis of the mutated genes variant allele frequency >10% showing the biological processes and pathways altered. **(E)** Percentage of mutated genes in each subtype of melanoma cells described in Tsoi et al., 2018.

## DISCUSSION

The signaling pathways that cooperate with MAPK/ERK activation in *BRAF*^*V600E*^ melanoma are of great interest. Several studies demonstrated that UVR is able to cooperate with the *BRAF*^*V600E*^ mutation, bypassing the oncogene-induced cell cycle arrest and promoting uncontrolled cell proliferation and tumor development (Luo et al., 2013; Viros et al., 2014). In addition to this, *Pten* or *Nf1* loss has been shown to abrogate *Braf*^*V600E*^-induced oncogene induced senescence (OIS) and lead to *in vivo* melanoma formation, and *Lkb1* loss has been shown abrogate *Braf*^*V600E*^-induced cell growth arrest without full progression to malignancy (Dankort et al., 2009; Gibney and Smalley, 2013; Vredeveld et al., 2012). Despite the well-known functions of LKB1 as a tumor suppressor, LKB1 also plays a relevant role in DDR (Esteve-Puig et al., 2014; Gupta et al., 2015; Ui et al., 2014; Wang et al., 2016), suggesting a cooperation between *BRAF*^*V600E*^ mutations, UVR-induced DNA damage and/or *LKB1* loss to promote melanomagenesis.

Initial analysis of human samples from the TCGA database supported previous observations suggesting the low expression or inactivation of *LKB1* in melanoma patients (Guldberg et al., 1999; Rowan et al., 1999). This data also showed a negative correlation between *BRAF* and *STK11* mRNA and protein expression. While many patients suffered shallow copy-number deletions in STK11 gene, BRAF locus was subjected to copy-number gains or amplifications. However, mRNA expression of *STK11* was more abundant than *BRAF*, which could be interpreted as a compensatory mechanism at transcriptional level that did not correlate with the corresponding amount of protein expression. This negative correlation of BRAF and STK11 at protein level was also observed in our validation subset of *BRAF*^*V600E*^-mutated human melanomas, supporting the notion of the lack of expression or low amounts of LKB1 in at least 50% of *BRAF*^*V600E*^ mutated samples. Due to this observation and the role played by LKB1 in DNA damage repair (Esteve-Puig et al., 2014; Gupta et al., 2015; Ui et al., 2014; Wang et al., 2016), we investigated the contributions of *Lkb1* loss to UVR-induced melanoma development and progression in a *Braf*^*V600E*^ mutational context. In contrast to previous report (Damsky et al., 2015), *Lkb1* haploinsufficiency cooperated with *Braf*^*V600E*^ promoting full tumorigenesis, although the progression occurred at a longer time than the previously reported. Further characterization of these tumors by immunohistochemistry also confirmed the activation of both pathways, mTORC1 and mTORC2/Akt for cell progression to malignancy (Damsky et al., 2015). *Lkb1* loss did not further increase *Braf*^*V600E*^ UVR-induced melanomas, probably due to the high tumor penetrance in the model. For unknown reasons currently under investigation, the loss of both *Lkb1* alleles *(Lkb1*^-/-^) promoted a delay in melanoma development. However, in agreement with the impaired UVB-induced DNA damage repair and the described role of LKB1 in DDR, the loss of *Lkb1*^-/-^ contributed to genetic instability increasing tumor multiplicity, even when compared with heterozygous mice (*Lkb1*^*+/-*^). This data also supports the acquisition of additional genetic alterations in the absence of UVB radiation, that will facilitate the promotion to full tumorigenesis of *Braf*^*V600E*^ melanocytes upon *Lkb1* loss.

Although three major histological morphologies were identified in most of the samples, melanomas with neural-like differentiation were more frequent upon *Lkb1* deletion. Tumor morphological heterogeneity increased upon *Lkb1* loss coincident with the genetic instability generated by the impaired DDR (Esteve-Puig et al., 2014; Gupta et al., 2015; Jin et al., 2020). The enrichment in neural-like morphology in *Lkb1* deficient tumors including the UV irradiated tumors, suggested a differential melanocyte subtype for tumor origin. This hypothesis was supported by the observed differential expression of LKB1 in melanocytes at seven days post-natal.

Mutational profiling of mouse tumor samples showed that 10% of the identified genes were also mutated in human samples, also showing a similar number of unique mutated genes supporting the relevance of the mouse model. UVB-irradiated samples accumulated C to T transitions while nonirradiated samples expression accumulated G to T transversions indicating the participation of alternative mutational mechanisms mostly in the absence of *Lkb1*. Nonsynonymous variants were the predominant type of mutation and stop gains were accumulated specially in irradiated samples. Non-irradiated *Lkb1* null samples showed a slight increase in non-frameshift deletions, showing also different rates of protein families mutated genes. *LKB1* deficiency increases the sensitivity of cells to radiation-induced carcinogenesis and ROS, resulting in excessive DNA oxidation and mutation rates (Gupta et al., 2015; Xu et al., 2015), affecting genomic stability in several ways and contributing to cancer development (Li and Zhu, 2020). Comprehensive analysis (Metascape) of the mutated genes lists according to the mouse genotype and treatment indicated that *Braf*^*V600E*^ tumors showed alterations in motility, adhesion, and cell signaling processes. Interestingly, *Lkb1* null tumors, independently of the UVB radiation status, showed a clear enrichment for neural differentiation related processes supporting the dedifferentiation and predominant neural-like morphology linked to those tumors. In fact, a number of publications have related LKB1 signaling with neural development and homeostasis (Kuwako and Okano, 2018), including the development of neural crest cell derivatives such as melanocytes (Radu et al., 2019). Due to the additional pleiotropic roles of LKB1 in cancer (i.e.: cell viability, invasiveness, and metabolism), the dysregulation of all these distinct aspects will also contribute to the mutation landscape selection and malignancy.

Collectively, this study and our previous reports (Esteve-Puig et al., 2009; Esteve-Puig et al., 2014; Gonzalez-Sanchez et al., 2013; Granado-Martinez et al., 2020) identify the loss of *LKB1* as an important mechanism cooperating with *BRAF*^*V600E*^ mutation in cancer development and progression. The loss of the multitask *LKB1* kinase will contribute to melanoma development through the dysregulation of multiple processes, including the increase of genomic instability caused by a deficient DDR. Loss of *LKB1*, not only makes cells especially vulnerable to UVB radiation and prone to cooperate with oncogenes and/or tumor suppressors, but promotes melanocyte transformation toward a neural-like phenotype. Thus, we identified the loss of *LKB1* as a mechanism cooperating with *BRAF*^*V600E*^ and UVB contributing to melanocyte transformation and melanomagenesis.

## Acknowledgements

This work was funded by Instituto de Salud Carlos III and co-funded by European Union (ERDF/ESF, “A way to make Europe”/ “Investing in your future”), PI17/00043-Fondos FEDER; PI20/0384-Fondos FEDER; PI23/00428-Fondos FEDER JAR, Euronanomed2-ISCIII (AC16/00019)-Fondos FEDER; JAR, Asociación Española Contra el Cancer (AECC-GCB15152978SOEN). AGAUR, 2021-SGR00653 JAR (supported PGM, KM);, Ramón Areces Foundation (supported KM and research); JAR.

## Author contributions

Conceptualization: JAR, KM. Investigation: KM, PG RO, EGS, BF JHL, IO, JMC, and JAR; Resources: JAR, VG and EMC; Funding acqusition: JAR, VGP. Methodology: KM, IO. Formal analysis: JAR and KM. Writing-review and editing: JAR, KM. Supervision: JAR

